# Application of a Latent Trait Modeling Method for Missing Data Across Datasets: Guidance on Appropriate Factor Structure

**DOI:** 10.1101/2022.11.14.516488

**Authors:** Christopher W. Bartlett, Tyler J. Gorham, Emily A. Knapp, Amii M. Kress, Brett G. Klamer, Steven Buyske, Bryan Lau, Stephen A. Petrill

## Abstract

Latent trait space can be leveraged to harmonize small data into big data when the constituent datasets measure the same underlying (latent) domains using a set of partially overlapping measurement instruments in each domain. The latent trait space then acts as a common metric space for each dataset, thus ensuring the same scale for the latent traits across datasets, despite the use of non-identical sets of measurement instruments within datasets. This approach, as originally published, only applied to a narrow set of circumstances, namely, that each measurement instrument occurred in more than one dataset. Here, we extend the latent trait approach to drop this requirement by using matrix completion methods. Using a simulation study, we evaluate the reliability of this extension and offer guidance on circumstances when the latent trait approach to missing data is robust and practical on real datasets.

---

The formation of large research consortia using collaborative study designs have gained popularity in recent years at the nexus of advances in data science and the need to create adequately powered datasets to improve replication (Fair, Dosenbach, Moore, Satterthwaite, & Milham, 2021). However, the “independent contributing studies” that participate in large research consortiums most often differ in study designs, protocols, and data measurement instruments (Lesko et al., 2018). Unique challenges in missing data arise in this context. In order to analyze big data from the consortium, common variables are needed but are often not available in all of the datasets.

The task of analyzing partially overlapping datasets lacks a mature model of best practices but has been evolving over time (Jacobson, Lau, Catellier, & Parker, 2018). For example, the consortium might perform a mega-analysis using variables that most commonly occur across the datasets. This approach greatly increases the sample size, and therefore power, over any single dataset but not leverage all of the datasets. Analytical approaches to this problem include, for example, combining the results of analyses in each datasets using metaanalysis (Haidich, 2010), and rescaling measures across the datasets for mega-analysis. Both approaches assume that combining measures across datasets that were not conducted with identical instruments or protocols has a reasonable interpretation phenotypically, and reasonable properties for statistical inference.

Use of meta-analysis, rescaling, or selective combining of datasets all have the virtue of being widely applicable regardless of the cross-dataset similarities and are both easy to calculate and intuitive to apply. However, consortium data often has a cross-dataset similarity at the measurement level that flows from the latent constructs that underpin the consortium. Leveraging this latent trait space is the alternative framework we present here that exploits the cross-dataset similarity structure commonly seen in collaborative design. The fundamental idea is sound *prima fascia*, namely, each constituent dataset in the consortium collected highly similar data across the same measurement domains else those datasets would not be a good fit to participate in the consortium. Each constituent dataset measured the similar latent traits, though with varying degrees of overlap in the specific measurement instrument used, but *en masse* the many of the same measurement instruments will be used in multiple studies and few instruments will be unique to one study. It is this cross-dataset similarity that a latent trait space approach can be successfully applied.

The first incarnation of this approach, or at least the first articulated attempt to use latent trait space for this problem, began with the strong simplifying assumption that the measures in the consortium appear in at least two constituent datasets. This assumption is valid as it will sometimes occur in collaborative designs, but often there will be one or more measurement instruments that that occur in a single dataset. The original formulation of this approach would simply ignore those incidental measures in the analysis. As presented in Figure 1, the grand correlation matrix across consortium data is complete when there are no incidential measures, but would be incomplete if any of these incidental measures are present. The latter renders the latent trait space incalculable.

**Figure 1.**
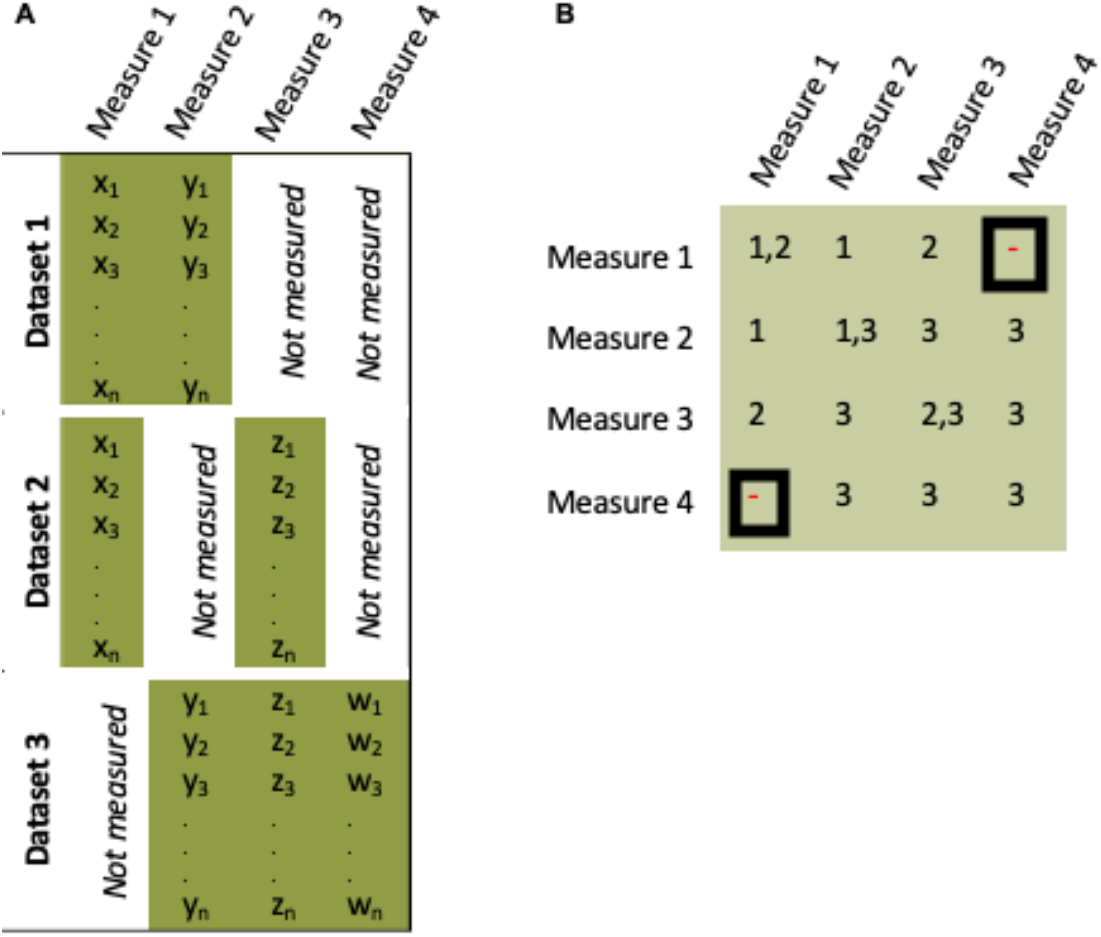
Minimal Example of How a “One-Off” Measurement Induces an Incomplete Correlation Matrix. In this set up, all four measures are correlated through a single unobserved latent trait (indicated by the green color background). A) Across the three datasets, there are no measurement instruments universally available. B) When calculated a correlation matrix across all datasets, the fact that Dataset 3 has a measure that occurs in no other dataset, plus the fact that Dataset 3 did is missing measurement 1, the grand correlation matrix is incomplete. An incomplete matrix precludes downstream factor analysis modeling.

As a motivating example of cross-dataset similarity, the authors on a large collaborative study design called the Environmental influence on Child Health Outcomes (ECHO) program. ECHO is a consortium of 69 pediatric cohort studies that focuses on modifiable aspects of the early life environment and its association with five outcome areas: perinatal health, obesity, respiratory disease, neurodevelopment, and positive health {LeWinn, 2021 #12}. ECHO represents a new model for collaborative research studies in that it is organized around a broad study population (child living in the US and Puerto Rico) rather than a specific exposure or disease outcome, and each cohort began with a unique focus and data collection protocol (Lesko et al., 2018). However, some measurement domains vital to the success of the program are common across the studies.

The prime example motivating the current research is socioeconomic status (SES) data collected under the 69 cohort-specific protocols, in addition to standardized data collection of SES for newly enrolled participants. In order to conduct pooled analysis across ECHO cohorts, differences in measurement, including cross-dataset similarity versus uniqueness must be taken into account. Socioeconomic status measures in ECHO protocol include income, supplementary information about the income such as household size supported by the income, sources of income, Supplemental Nutrition Assistance Program and WIC program benefits, housing, measures of economic insecurity, detailed employment information, and the level of education attained by the mother and father. Prior to adopting the ECHO-wide protocol, cohorts collected socioeconomic status in a variety of ways. Previous harmonization efforts were undertaken to combine as much data across cohorts as possible. ECHO cohorts represent heterogenous populations in terms of many factors related to SES: race/ethnicity, geography; and some cohort selection factors are direct measures of SES: individual income, low-income residential areas, or recruitment from clinics serving low-income populations.

The present research was motivated by both the appeal of applying the latent trait approach to the subdomains of SES and the need to address the “one off” measurement problem in the epidemiological collaborative study design used by ECHO. In this practical setting, we sought to address the incomplete grand correlation matrix problem and also to take the first steps in developing practical guidance on when the latent trait space approach to mega-analysis is reasonable to apply, in the context of ECHO but also more generally for the wider behavioral and epidemiological research communities. In the sections that follow, we explain the latent trait space method we initially developed as an R library called *Rosetta* (Bartlett, Klamer, Buyske, Petrill, & Ray, 2019). We then describe a numerical problem that arise in collaborative studies where the grand correlation matrix can be incomplete in some circumstances. We then discuss the matrix completion algorithm that can mitigate the incomplete matrix problem. To assess the matrix completion algorithm in the context of combining data for collaborative designs, we discuss a simulation study we conducted.

## The Latent Trait Space Approach

The latent trait space approach solves a pattern of missing data across the constituent datasets that often occurs in consortia. This pattern can be readily observed in simplified example using only one latent trait domain (Figure 2). Within a measurement domain, each dataset has at most two measurement instruments for that domain. Consider a consortium where, in total, across three datasets there are three measurement instruments but within any given constituent dataset, only two of the three measurement instruments were used. Given this scenario we can deduce the following. 1) A mega-analysis of any single measurement will necessarily result in missing data from at least one of constituent datasets. 2) A meta-analysis could be used to combine p-values across datasets but this would require making the strong assumption (from a measurement point of view) that agglomerating analysis results across different measurement instruments is valid and interpretable. 3) A mega-analysis of data that has been rescaled within dataset makes highly similar assumptions as a meta-analysis with regard to the validity and interpretability of the rescaling operation.

**Figure 2.**
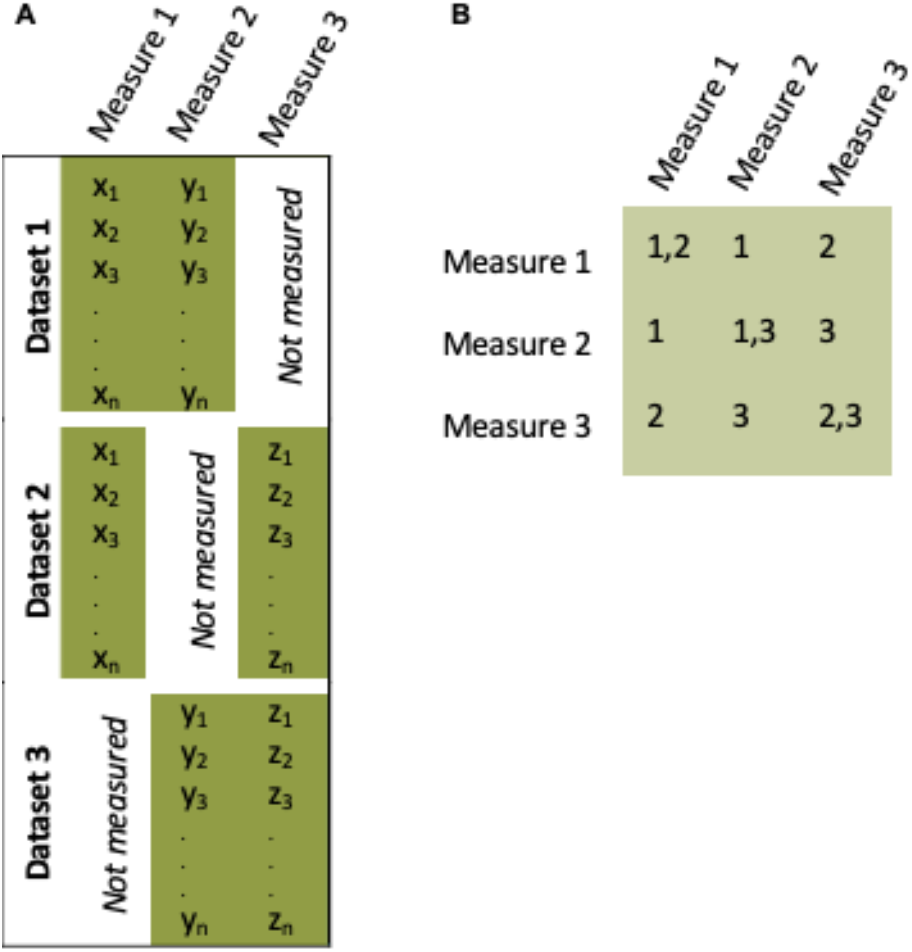
A Simplified Collaborative Study Design Where No Measurements Are Universally Available. All three measures are correlated through a single unobserved latent trait (indicated by the green color background). A) Across the three datasets, there are no measurement instruments universally available, thus precluding a sampling pooling of the data for mega-analysis. B) Despite the missing data, a grand correlation matrix can be calculated completely.

Latent trait models have a set of assumptions that directly bear on the given analysis scenario. Deriving the latent trait is a well appreciated method to reduce measurement error in analysis and has a long and distinguished history in psychometrics and the behavioral sciences more generally (Schmitt, 2011). The latent trait space approach to analysis of collaborative study designs inherits this important property. In particular, the measurement instruments should be correlated through a common latent factor. Statistically, we state that after conditioning on the latent trait those instruments would be uncorrelated as the common variance is completely accounted for by the underlying latent trait. It follows that each measurement instrument has both common variance with the underlying latent trait, but that random measurement error is uncorrelated across the instruments.

The standard statistical assumptions for linear modeling also apply to latent trait analysis. Chiefly among these is the Gaussian distribution of the latent trait itself. Latent trait are not directly observable, but as a model of a continuous trait across the populations of interest, this is largely true in behavioral domains of interest and much of epidemiology as well. Note the observed data need not be Gaussian. The assumption of linearity is quite strong. Here we note linearity in terms of the function that maps data to the latent trait space and not the shape of the data per se. That is to say, we expect that the mapping function have the following two properties: 1) *f*(*x* + *y*) = *f*(*x*) + *f*(*y*) and, 2) *f*(*αx*) = *αf*(*x*). The standard equation of a line (*f*(*x*) ≡ *mx* + *b*) satisfies the two conditions of linearity. Presenting the two underlying assumptions of linearity is not meant to obscure the assumption, but instead will make a later step in the *Rosetta* calculation easier to understand.

Considering the simplified scenario in Figure 2, the latent trait approach to this pattern of missing data is to create one domain score (the latent trait score) that represents the same underlying latent trait, which necessarily has the scale (i.e., the same mean and variance). Deriving the latent trait solves the missing data problem without subsetting the data (dropping datasets from the analysis) and the derivation of the latent trait combines the data across datasets without ad hoc assumptions needed by meta-analysis or manual rescaling. Furthermore, latent traits are without random measurement error. For these reasons, when the structure of the data is appropriate, the latent trait space approach will increase statistical power over the alternatives. When multiple correlated measurement domains are included in the model, the accuracy of latent trait estimation will improve as the correlated traits provide additional information for estimating each latent trait and offer further reduction in measurement error. Yet, there are two remaining problems that limit the application of this approach to more complex (and common) missing data structures in consortia: The One-Off Problem and the Matrix Completion Problem

### The One-Off Problem

To define why the One-Off Problem arises, the simplified scenario in Figure 2 needs to be expanded. For one, the datasets needs to include a measurement instrument that does not occur in the other two datasets (like Figure 1). Contrasted with Figure 2, a missing measurement in a dataset, even if every dataset has missing measurements, the grand correlation matrix can still be completely defined. The situation where the grand correlation matrix is not complete occurs when one dataset has a unique measure and also has at least one other measurement instrument missing. Thinking about the correlations pairwise, it is not possible to calculate the correlation between the unique measurement instrument and any instrument not measured in that dataset. Hence, the grand correlation matrix will have missing information. Much more complicated scenarios for inducing missing data in the grand correlation matrix are possible, but we present the “one-off” problem as the minimally defined situation where this occurs. Solving this problem generalizes to other more complex scenarios.

Moreover, as mentioned above, the latent trait space approach is ideally applied when there are multiple correlated latent traits is theoretically and empirically justified. By expanding both the number of measurement instruments in one dataset and expanding the number of measurement domains (i.e., the number of latent traits), we have a simple but realistic scenario that offers the essence of consortium data analysis problem and is still being wieldy for simulation studies and discussion (presented in Figure 3). In the simulation study, below, we also present two variations of the scenario in Figure 3 that increase the complexity of the missing data.

**Figure 3.**
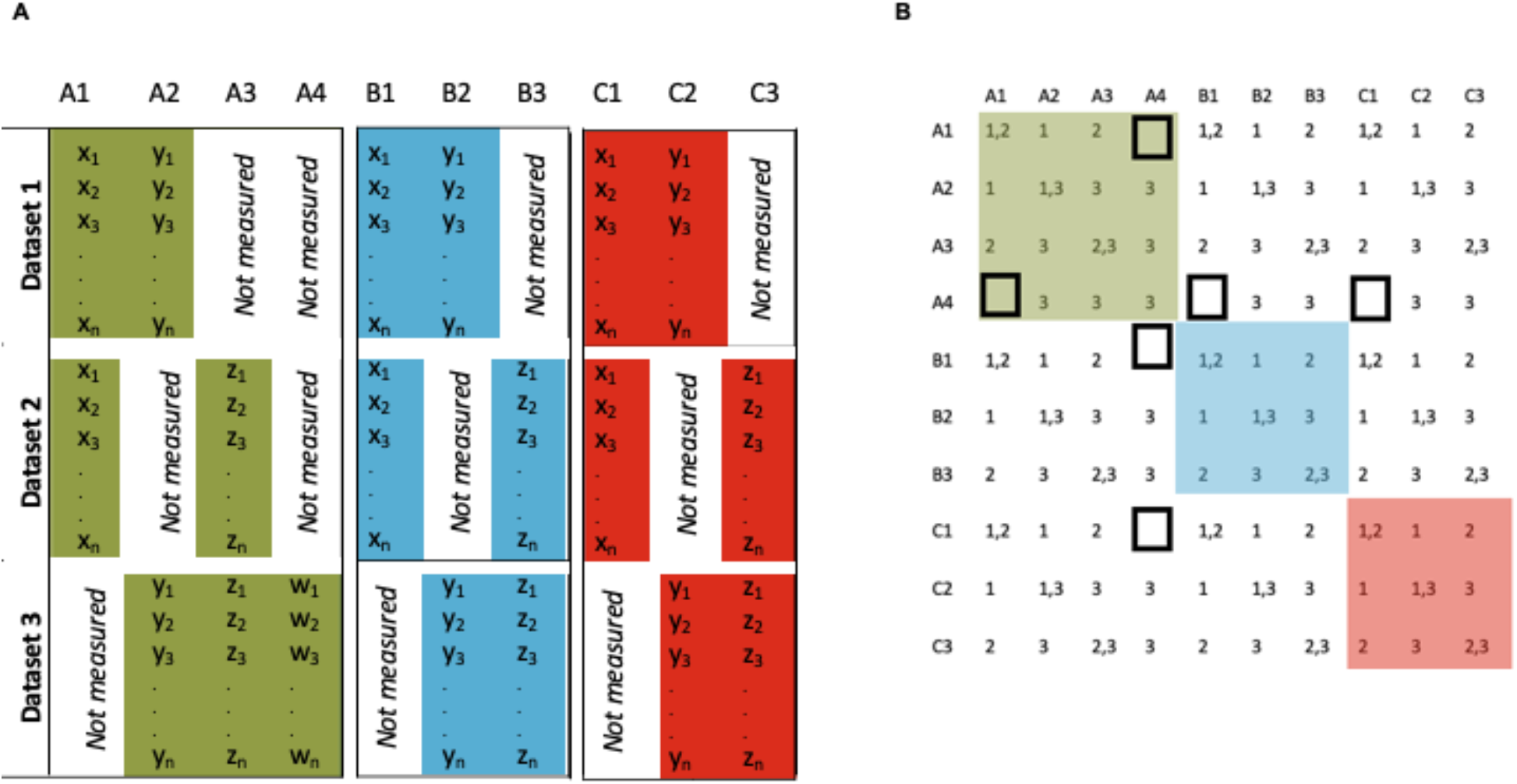
A Realistic Collaborative Study Design With an Incomplete Correlation Matrix. Measurements are correlated through a set of three unobserved latent traits (indicated by the green, blue, and red color backgrounds, respectively). A) Across the three datasets, there are no measurement instruments universally available in each domain, thus precluding a sampling pooling of the data for mega-analysis. B) Despite the missing data, a grand correlation matrix can be calculated with only 6 missing entries, that are induced by the “one-off” measurement whereby Dataset 3 is the only dataset with measurement A4.

### Matrix Completion Problem

Numerical linear algebra has long studied the matrix completion problem, defined as finding values that complete missing information in a matrix given a set of constraints (such as requiring only integers, only positive values, sparsity requirements, etc). Here the constraints are quite favorable. A correlation matrix has diagonal values all equal to 1, is symmetric, and is positive semidefinite (all eigenvalues ≥ 0). Previous work shows that first finding missing correlation values that maximize the determinant (MaxDet) of the correlation matrix (Georgescu, Higham, & Peters, 2018), and second shrinkage of the resultant matrix (adjusted through scalar multiplication) provides the nearest correlation matrix from the estimated values under the MaxDet procedure (Higham, 2002).

## Goals of the Study

Numerical linear algebra routines can be difficult to work with in real data situations. Analysts with practical experience in factor analysis or any latent trait method has surely received warnings and errors when processing real data. Thus while we know of methods for matrix completion that should solve the “one-off” problem for our application, empirical study is needed to assess the robustness of any claim. In the simulation study described in the Methods section, we outline our approach to systematic testing of the robustness of matrix completion recently implemented in the *Rosetta* R library, our implementation of the latent trait space approach.

As mentioned earlier in the introduction, the widely used approaches to the analysis of collaborative study designs are applicable to a very broad range of datasets and data structures. The latent trait space approach is more limited in the scope of utility and we set out to define guidance on when our approach is expected to be applicable. Given that latent traits are defined by correlations within and between measurement domains, we hypothesized that once the within and/or between domain correlations are too low the method lacks utility and should not be applied. Our simulation study was designed to find those thresholds to provide a reasonable guide for future studies.

## Methods

In a simulation study, datasets are created using random numbers generated to fit a prespecified generating model. Simulation studies are helpful when studying numerical routines and proposed statistical methods since an underlying true model can be used for comparisons to assess error variance from a statistical model. Results from real datasets do not have a ground truth for comparison, so even though the ECHO program was the motivation for this study, we do not apply the method to ECHO data. The purpose of this simulation study is validation of the matrix completion method and to define guidance on when *Rosetta* is expected to have advantages over other methods for combining datasets.

### The Rosetta Algorithm

The *Rosetta* algorithm implements the latent trait space approach and has been described previously. Briefly, a grand correlation matrix over all datasets is built using logical intersections of the datasets for each pairwise correlation to ensure each calculated correlation uses the maximum possible number of datasets with two measurements needed for a given correlation. Eigen decomposition of the matrix is performed to estimate eigenvalues and eigenvectors which are then normalized to factor loadings and communalities. The number of factors can be estimated from the data or specified by the user. Using a structural equation framework, correlations within factors are constrained to the grand correlation matrix (similar to confirmatory factor analysis) and correlations between factors are also constrained by the previous step (not done in confirmatory factor analysis). These constraints ensure that the factor scores output by Rosetta are on the same scale across datasets, since by the linearity assumption, combinations of constituent measures lie within the same latent trait space.

### Simulations

The generating model for simulations is based on four grand correlation matrices that correspond to four scenarios:

I. Baseline model of three measurement domains (A, B and C), each with three measurement instruments (A1, A2, A3, B1, B2, B2, C1, C2, C3)
II. An extension of model I with an added “one-off” measure in domain C (measure C4 added to only one constituent dataset)
III. An extension of model II with another “one-off” measure added in domain B (measure B4 added to a different constituent dataset than C4)
IV. An extension of model I with two additional “one-off” measures, both added to the same dataset (C4, C5)

These four models offer a baseline for comparisons (model I) that has a complete matrix versus three scenarios where matrix completion is necessary (models II, III, and IV). The main comparison of interest is between models I and II to assess if our matrix completion is adequate in the minimally deficient scenario. The remaining models will assess how robust the matrix completion is to increasing levels of missing information. Each model requires specification of a grand correlation matrix as input into the simulation (see Supplemental Tables).

Within-domain correlations were varied in the set [0.1, 0.2, 0.3, 0.4, 0.5, 0.6, 0.7, 0.8, 0.9] and between-domain correlations were varied in the set [0.1, 0.2, 0.3, 0.4, 0.5]. While the lowest (highest) correlations are lower (higher) than what is reasonable for real datasets, the extremes offer information on when the method would be useful and what the best possible performance that could be attained could be when planning a study. Correlations were the same between measures, and the same between domains equal to the simulated parameter. While this constraint is not realistic in practice, intuitively our simulation represents the average correlations that might be observed in real datasets and provides suitable representation on that basis.

Once data are simulated, we expect that data from the simulation will provide sample parameter estimates close to the population parameters from the simulation generating model. Within each model, we simulate a control condition that has complete data (i.e., no matrix completion necessary) by simulating all measurements in all datasets. Analysis of complete data are called “complete” runs where the same dataset with measures missing in multiple datasets is created by dropping data from the complete dataset. Analyses performed on the dataset with missing information is referred to as a “Rosetta” run. Correlations of approximately 1 between the “complete” and “Rosetta” runs would indicate optimal matrix completion.

## Results

For Scenario A, we observed highly similar patterns to the previous study, namely, that Rosetta versus complete data preforms nearly identically with better performance for higher correlations. Scenario B is the main situation of interest as it tests not only Rosetta with realistic complexity but also the matrix completion routine. The matrix completion routine works better as the correlation is constrained (higher correlations across measures and domains) seen in Figure 4. Performance in Scenarios C and D perform similar to Scenario B, showing that the matrix completion scales with the number of missing entries in the grand correlation matrix.

**Figure 4.**
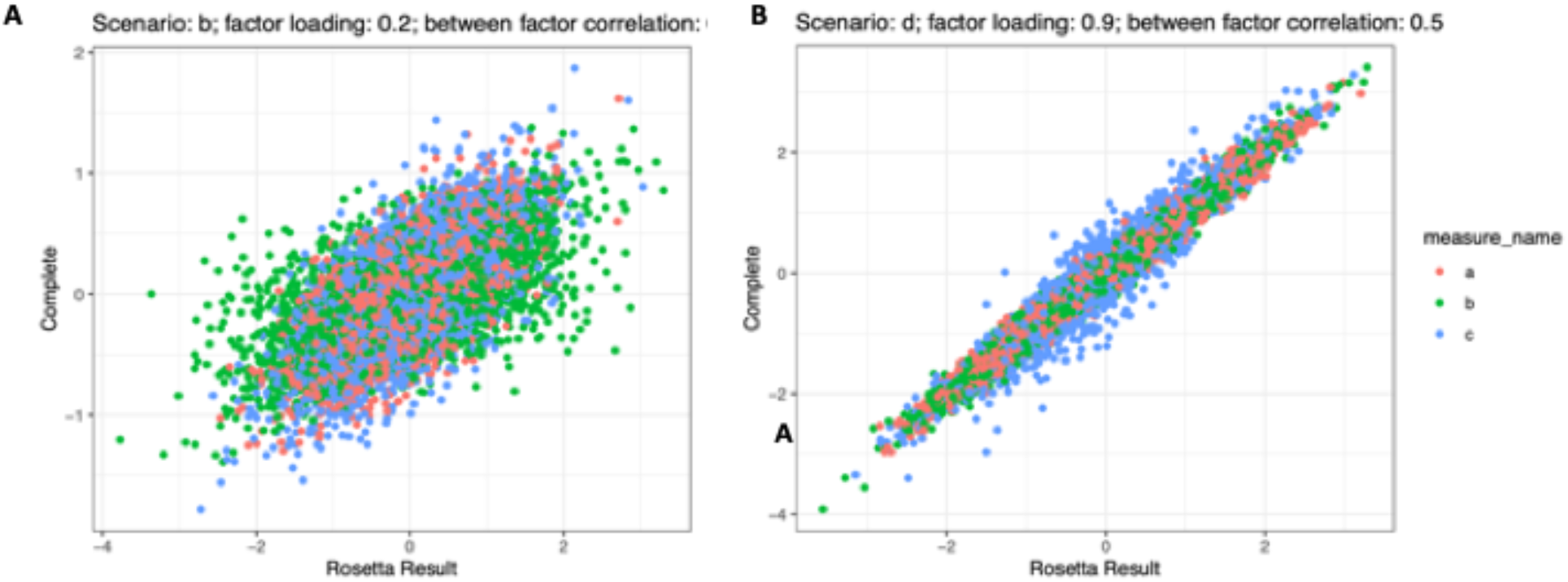
Example Plots for *Rosetta* Performance Under High and Low Correlation Conditions. We compared values of the latent traits under complete data versus Rosetta output for two conditions, a low correlation condition (0.2 within measures and 0.2 between latent trait domains) and a high correlation condition (0.9 within measures and 0.5 between latent trait domains).

### Numerical Stability

When the simulated correlations are low, numerical instability for factor analytic methods are expected. To quantify this effect, we simulated data and applied Rosetta and calculated the proportion of runs where numerical instability caused a failed exit status of the function (i.e., the run failed). The overall trend is seen in the heatmap in Figure 5. The chance of success is greater when both the within and between correlations are higher, as expected, but the success rate for conditions as low as 0.1 within correlation and 0.1 between correlations can be as high 90% for some data configurations.

**Figure 5.**
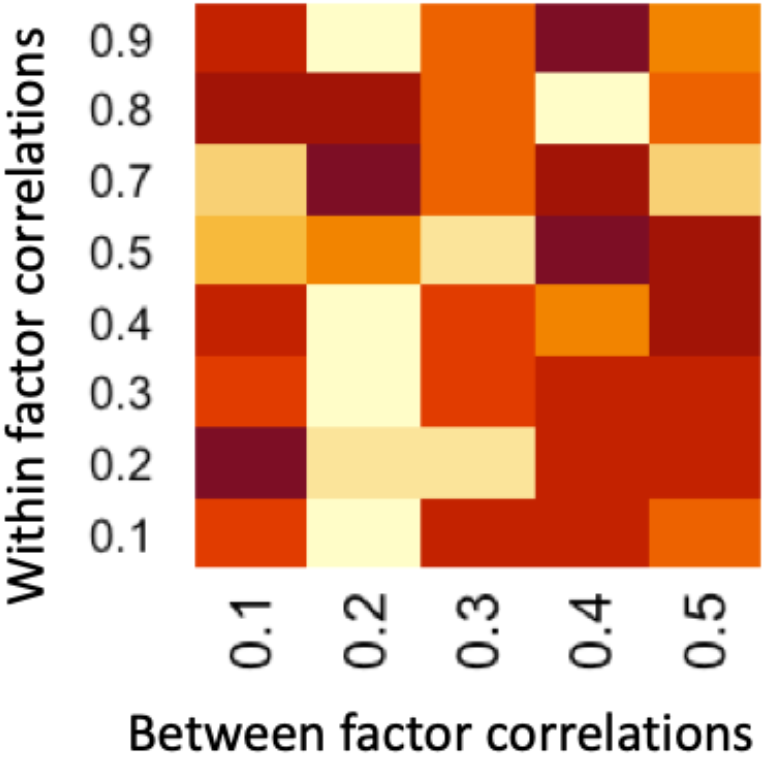
Numerical Success Rate by Within/Between Factor Correlations. A general trend for higher success rates with increasing correlations is observed.

## Discussion

The big data revolution has provided tools for ever increasing amounts of data but the question of heterogeneous data structures within a big dataset is still being developed. While successful and widely deployable tools already exist for combining datasets, such as metaanalysis and rescaling, we studied a recently developed approach that exploits a data structure commonly seen in collaborative designs. The recently developed latent trait approach implemented in *Rosetta* decreases measurement error variance and puts all constituent datasets on the same latent trait scale for mega-analysis. Mega-analysis of all datasets offers researchers the flexibility with modeling across an entire consortium.

The matrix completion method in Rosetta performs well for the parameters studied. When the simulated correlations are low, numerical instability for factor analytic methods are expected. In this study we sought to find a practical guideline for when Rosetta might be reasonably applied, when the correlation structure is not so low as to greatly reduce the chance of success. The reasons for numerical failures we observed included statuses such as ill-conditioned matrix, singular matrix, and too many iterations causing failed convergence. Not all of these issues need preclude a successful analysis for the experienced practitioner, but most would be difficult or impossible for an analyst not thoroughly experienced in latent trait analysis. While our method for tracking failure is therefore conservative, it is likely to be a good approximation of success rates for many end users. The complication is that the presence of numerical instability is reasonably stochastic. Even with correlations as low as 0.1 the success rate can be as high as 90%. Further work is needed to automate remediation for errors that can be corrected. Overall though, the trend we observed fits well with the general practice that a study with low correlations overall is not a good candidate for latent trait modeling based on empirical considerations alone.

All studies have limitations, and while simulation studies are easy to conduct, the set of parameter values evaluated is necessarily limited in scope. Here we presented scenarios that do occur in real datasets utilizing a collaborative study design and also tested a wider range of parameters than would be reasonable to observe in practice. Jointly these conditions offer a degree of ecological validity to our evaluations but cannot be taken as mathematical proof. And whenever numerical routines are so critically involved, care must always be taken in applying a method.

With regard to our goal of providing practical guidance on when to use *Rosetta*, it appears that the standard guidance on all latent trait modelling applies here. When the correlation structure strongly indicates that latent traits may be at play and there are additional theoretical and empirical grounds to further justify their use, then Rosetta might be productively attempted. Should numerical instability be an issue, then the standard methods are always available in those instances.

## Article Information

### Ethical principles

The authors affirm having followed professional ethical guidelines in preparing this work. These guidelines include obtaining informed consent from human participants, maintaining ethical treatment and respect for the rights of human or animal participants, and ensuring the privacy of participants and their data, such as ensuring that individual participants cannot be identified in reported results or from publicly available original or archival data.

## Acknowledgements

The authors would like to thank the families enrolled in the Environmental influences of Child Health Outcomes (ECHO) who provided the inspiration to understand how to best combine many smaller datasets into a large dataset to improve public health research

## Supplemental Tables

**Supplemental Table 1.**
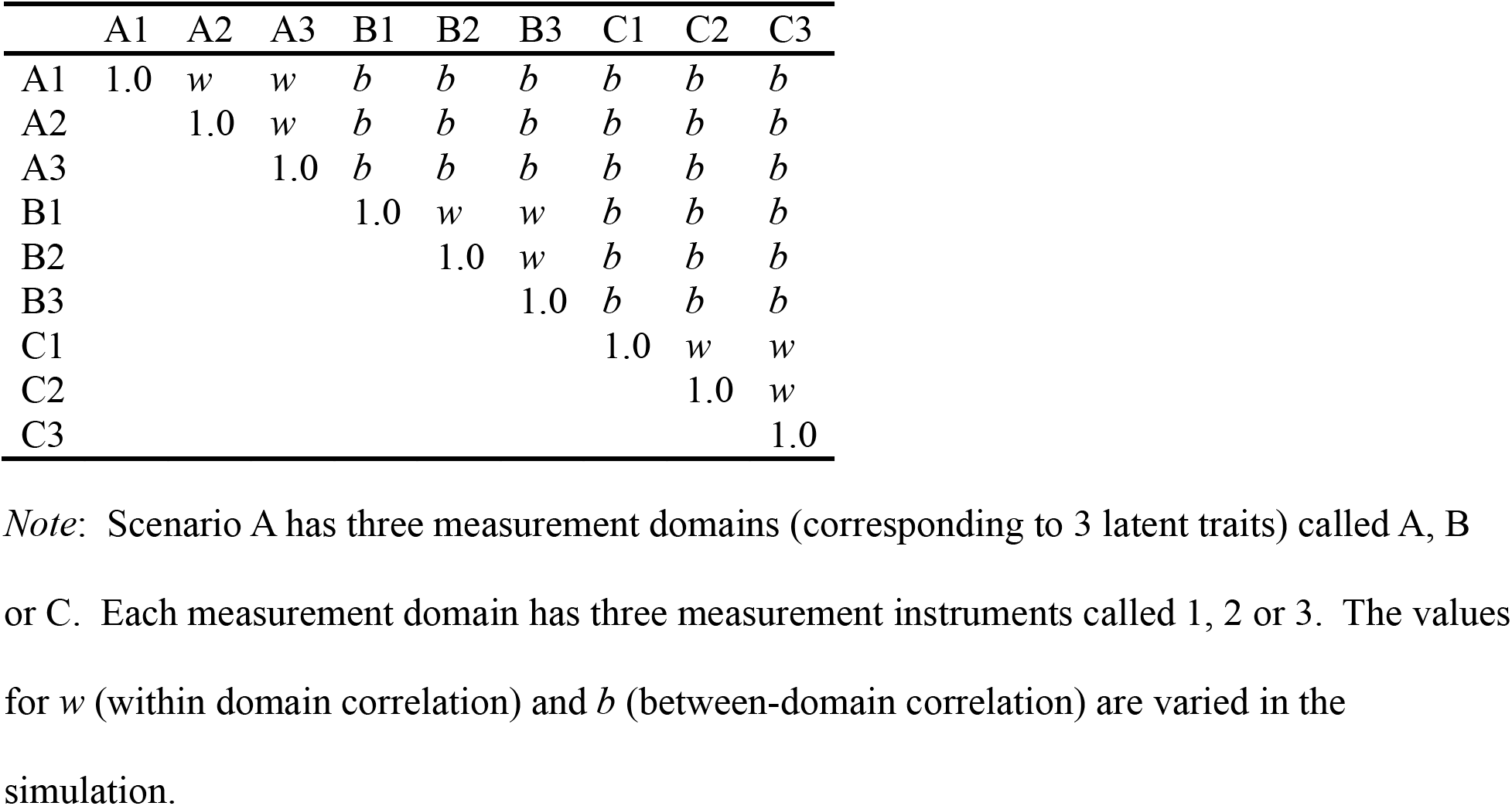
Correlation Matrix for Generating Condition: Scenario A

**Supplemental Table 2.**
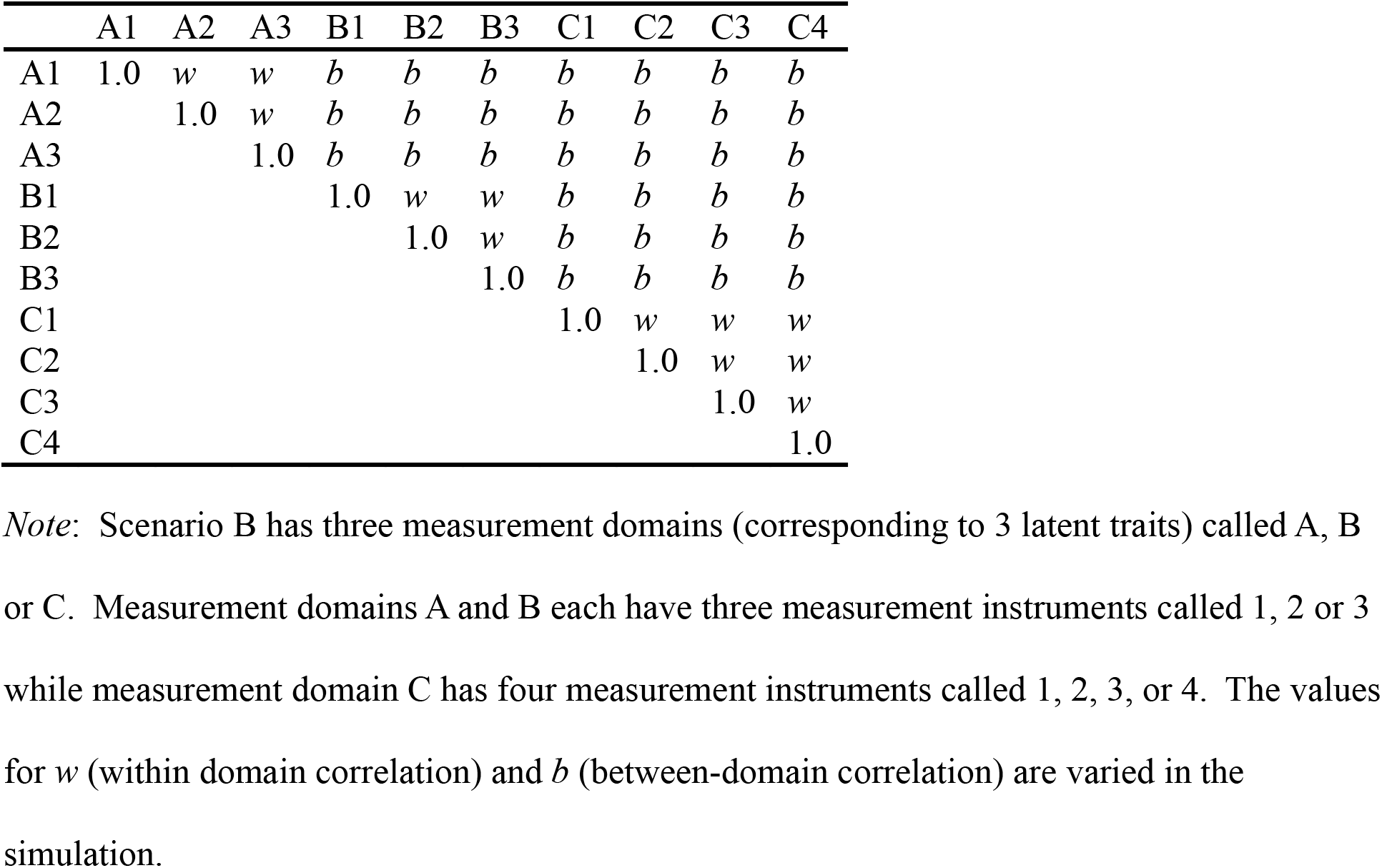
Correlation Matrix for Generating Condition: Scenario B

|  | A1 | A2 | A3 | B1 | B2 | B3 | C1 | C2 | C3 | C4 |
| --- | --- | --- | --- | --- | --- | --- | --- | --- | --- | --- |
| A1 | 1.0 | *w* | *w* | *b* | *b* | *b* | *b* | *b* | *b* | *b* |
| A2 |  | 1.0 | *w* | *b* | *b* | *b* | *b* | *b* | *b* | *b* |
| A3 |  |  | 1.0 | *b* | *b* | *b* | *b* | *b* | *b* | *b* |
| B1 |  |  |  | 1.0 | *w* | *w* | *b* | *b* | *b* | *b* |
| B2 |  |  |  |  | 1.0 | *w* | *b* | *b* | *b* | *b* |
| B3 |  |  |  |  |  | 1.0 | *b* | *b* | *b* | *b* |
| C1 |  |  |  |  |  |  | 1.0 | *w* | *w* | *w* |
| C2 |  |  |  |  |  |  |  | 1.0 | *w* | *w* |
| C3 |  |  |  |  |  |  |  |  | 1.0 | *w* |
| C4 |  |  |  |  |  |  |  |  |  | 1.0 |

*Note:* Scenario B has three measurement domains (corresponding to 3 latent traits) called A, B or C. Measurement domains A and B each have three measurement instruments called 1, 2 or 3 while measurement domain C has four measurement instruments called 1, 2, 3, or 4. The values for *w* (within domain correlation) and *b* (between-domain correlation) are varied in the simulation.

**Supplemental Table 3.**
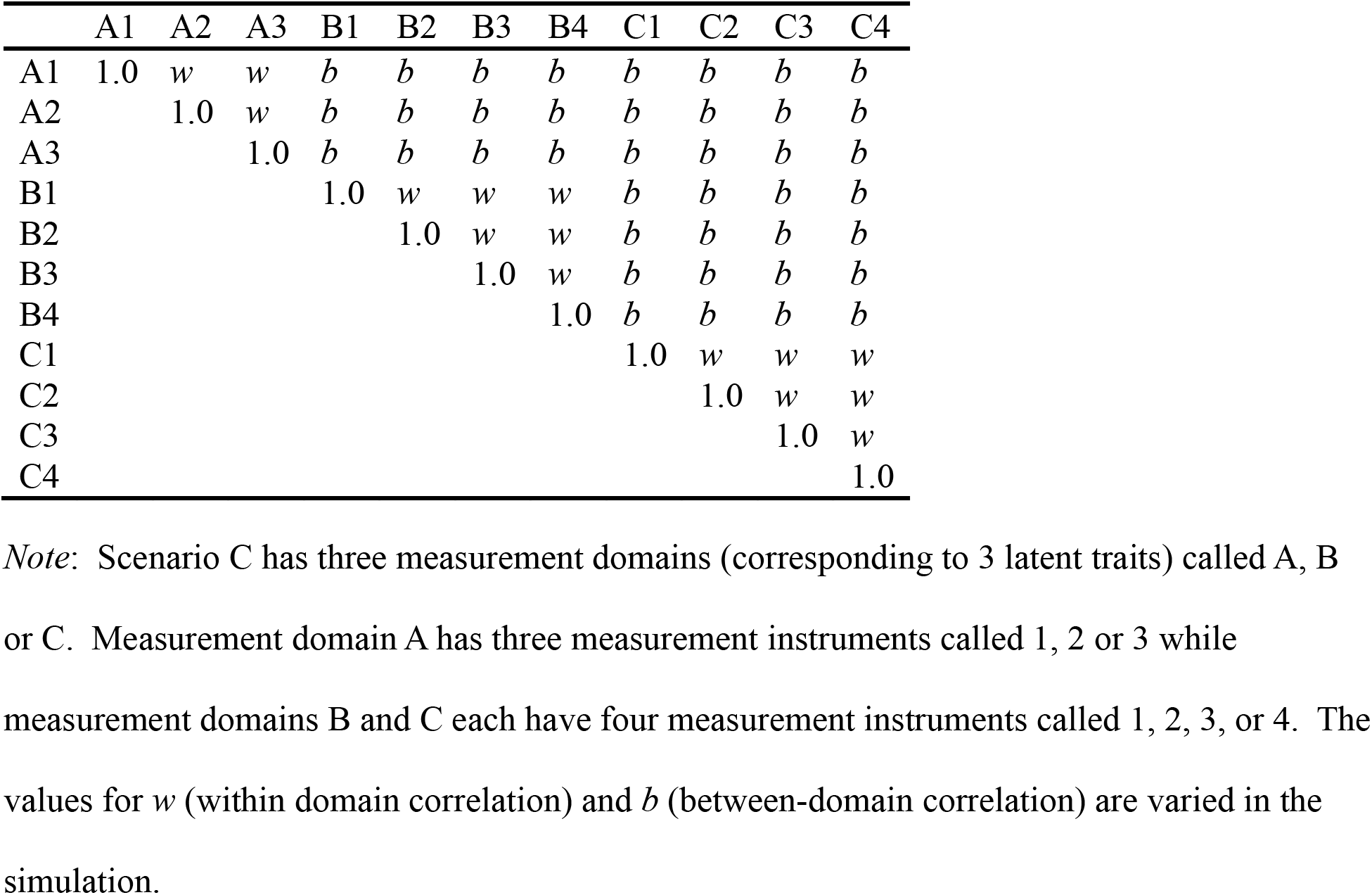
Correlation Matrix for Generating Condition: Scenario C

|  | A1 | A2 | A3 | B1 | B2 | B3 | B4 | C1 | C2 | C3 | C4 |
| --- | --- | --- | --- | --- | --- | --- | --- | --- | --- | --- | --- |
| A1 | 1.0 | *w* | *w* | *b* | *b* | *b* | *b* | *b* | *b* | *b* | *b* |
| A2 |  | 1.0 | *w* | *b* | *b* | *b* | *b* | *b* | *b* | *b* | *b* |
| A3 |  |  | 1.0 | *b* | *b* | *b* | *b* | *b* | *b* | *b* | *b* |
| B1 |  |  |  | 1.0 | *w* | *w* | *w* | *b* | *b* | *b* | *b* |
| B2 |  |  |  |  | 1.0 | *w* | *w* | *b* | *b* | *b* | *b* |
| B3 |  |  |  |  |  | 1.0 | *w* | *b* | *b* | *b* | *b* |
| B4 |  |  |  |  |  |  | 1.0 | *b* | *b* | *b* | *b* |
| C1 |  |  |  |  |  |  |  | 1.0 | *w* | *w* | *w* |
| C2 |  |  |  |  |  |  |  |  | 1.0 | *w* | *w* |
| C3 |  |  |  |  |  |  |  |  |  | 1.0 | *w* |
| C4 |  |  |  |  |  |  |  |  |  |  | 1.0 |

*Note:* Scenario C has three measurement domains (corresponding to 3 latent traits) called A, B or C. Measurement domain A has three measurement instruments called 1, 2 or 3 while measurement domains B and C each have four measurement instruments called 1, 2, 3, or 4. The values for *w* (within domain correlation) and *b* (between-domain correlation) are varied in the simulation.

**Supplemental Table 4.** Correlation Matrix for Generating Condition: Scenario D

|  | A1 | A2 | A3 | B1 | B2 | B3 | C1 | C2 | C3 | C4 | C5 |
| --- | --- | --- | --- | --- | --- | --- | --- | --- | --- | --- | --- |
| A1 | 1.0 | *w* | *w* | *b* | *b* | *b* | *b* | *b* | *b* | *b* | *b* |
| A2 |  | 1.0 | *w* | *b* | *b* | *b* | *b* | *b* | *b* | *b* | *b* |
| A3 |  |  | 1.0 | *b* | *b* | *b* | *b* | *b* | *b* | *b* | *b* |
| B1 |  |  |  | 1.0 | *w* | *w* | *b* | *b* | *b* | *b* | *b* |
| B2 |  |  |  |  | 1.0 | *w* | *b* | *b* | *b* | *b* | *b* |
| B3 |  |  |  |  |  | 1.0 | *b* | *b* | *b* | *b* | *b* |
| C1 |  |  |  |  |  |  | 1.0 | *w* | *w* | *w* | *w* |
| C2 |  |  |  |  |  |  |  | 1.0 | *w* | *w* | *w* |
| C3 |  |  |  |  |  |  |  |  | 1.0 | *w* | *w* |
| C4 |  |  |  |  |  |  |  |  |  | 1.0 | *w* |
| C5 |  |  |  |  |  |  |  |  |  |  | 1.0 |

*Note:* Scenario D has three measurement domains (corresponding to 3 latent traits) called A, B or C. Measurement domains A and B each have three measurement instruments called 1, 2 or 3. Measurement domain C has five measurement instruments called 1, 2, 3, 4, or 5. The values for *w* (within domain correlation) and *b* (between-domain correlation) are varied in the simulation.

## References

Bartlett, C. W., Klamer, B. G., Buyske, S., Petrill, S. A., & Ray, W. C. (2019). Forming Big Datasets through Latent Class Concatenation of Imperfectly Matched Databases Features. Genes (Basel), 10(9). doi:10.3390/genes10090727

Fair, D. A., Dosenbach, N. U., Moore, A. H., Satterthwaite, T. D., & Milham, M. P. (2021). Developmental cognitive neuroscience in the era of networks and big data: Strengths, weaknesses, opportunities, and threats. Annual Review of Developmental Psychology, 3, 249–275.

Georgescu, D. I., Higham, N. J., & Peters, G. W. (2018). Explicit solutions to correlation matrix completion problems, with an application to risk management and insurance. R Soc Open Sci, 5(3), 172348. doi:10.1098/rsos.172348

Haidich, A. B. (2010). Meta-analysis in medical research. Hippokratia, 14(Suppl 1), 29–37. Retrieved from http://www.ncbi.nlm.nih.gov/pubmed/21487488

Higham, N. J. (2002). Computing the nearest correlation matrix—a problem from finance. IMA journal of Numerical Analysis, 22(3), 329–343.

Jacobson, L. P., Lau, B., Catellier, D., & Parker, C. B. (2018). An Environmental influences on Child Health Outcomes viewpoint of data analysis centers for collaborative study designs. Curr Opin Pediatr, 30(2), 269–275. doi:10.1097/MOP.0000000000000602

Lesko, C. R., Jacobson, L. P., Althoff, K. N., Abraham, A. G., Gange, S. J., Moore, R. D., … Lau, B. (2018). Collaborative, pooled and harmonized study designs for epidemiologic research: challenges and opportunities. Int J Epidemiol, 47(2), 654–668. doi:10.1093/ije/dyx283

Schmitt, T. A. (2011). Current methodological considerations in exploratory and confirmatory factor analysis. Journal of Psychoeducational Assessment,, 29(4), 304–321.

